# Habitat Suitability modeling of Urial (Ovis orientalis arkal) in the Samelghan plain by using Maximum Entropy Method

**DOI:** 10.1101/2021.11.29.470339

**Authors:** Abbas Naqibzadeh, Jalil Sarhangzadeh, Ahad Sotoudeh, Marjan Mashkur, Judith Thomalsky

## Abstract

Habitat suitability models are useful tools for a variety of wildlife management objectives. Distributions of wildlife species can be predicted for geographical areas that have not been extensively surveyed. The basis of these models’ work is to minimize the relationship between species distribution and biotic and abiotic environments. For some species, there is information about presence and absence that allows the use of a variety of standard statistical methods, however, the absence data is not available for most species. Nowadays, the methods that need presence-only data are expanded. One of these methods is the Maximum Entropy (MaxEnt) modeling. The purpose of this study is to model the habitat of Urial (*Ovis orientalis arkal*) in the Samelghan plain in the North East of Iran with the MaxEnt method. This algorithm uses the Jackknife plot and percent contribution values to determine the significance of the variables. The results showed that variables such as southern aspects, Juniperus-Acer, Artemisia-Perennial plants, slope 0-5%, and asphalt road were the most important factors affecting the species’ habitat selection. The area under curve (AUC) Receiver Operating Characteristic (ROC) showed an excellent model performance. Suitable habitat was classified based on the threshold value (0.0513) and the ROC, which based on the results 28% of the area was a suitable habitat for Urial.

## Introduction

Suitable habitat has a significant impact on the survival and reproduction of species, management, and conservation of wildlife [1,2]. Global diversity in recent decades has declined due to land-use changes, climate change [3], habitat destruction and fragmentation, invasive species, overexploitation [4].

Habitats modeling is predicting species geographical distribution based on environmental conditions of known sites, as well as an important method in analytical biology with applications in environmental protection and planning, evolution, epidemiology, ecology, management of invasive species, and other fields [5]. Species distribution models (SDMs) are empirical models relating field observations to environmental predictor variables based on statistically or theoretically derived response surfaces [6,2]. SDMs estimate the relationship between species records at sites and the environmental and/or spatial characteristics of those sites [7,2,8].

Generally, due to the irregular distribution of species in habitats, it is difficult and costly to determine the exact distribution of species [9], therefore, such models are naturally static and probable, because statistically, the geographical distribution of species or communities are related to their current habitats [10], so these models can perform well in describing the natural distribution of species (within their current range). It is essential to accurately estimate the distribution of species, to ensure that environmental protection planning efforts are more beneficial in managed areas [9]. Species distribution models or habitat suitability models allow potentially predicting human effects on biodiversity patterns at different spatial scales, the origin more of modeling methods to predict of fauna and flora distribution have in environmental-species relations [11]. Habitat models of habitat-wildlife relationships are used to assess the potential of the area [12] and to create habitat patterns for species that are introduced or present in the area [13,14]. Digital distribution maps of species for basic and applied environmental research [15] based on powerful statistical software and Geographic Information System (GIS) tools, depending on the environmental needs of the species and their geographical distribution leads to the development of habitat modeling [16]. For some species, there is information about presence and absence that allows the use of a variety of standard statistical methods, however, the absence data is not available for most species [17].

Habitat evaluation models are divided into two groups, the group that needs presence and absence data and the group that needs the presence-only data, achieving the right non-presence data requires continuous monitoring of the habitat, recording the presence and absence of species for many years, and obtaining sufficient information about species ecology [18], because, reliable information about the absence of wildlife species cannot be easily obtained due to elusive behavior and patterns of activity [19], methods based on presence and absence data are exposed to the phenomenon of pseudo-absence. In other words, species observation by the observer for a variety of reasons, such as observer accuracy, equipment used, species behavior in camouflage, and concealment, causes that point to be recorded as the non-presence. This can lead to errors in data analysis [20]. Therefore, the use of models that only require the presence can prevent using pseudo-absence data [21]. As a result, modeling techniques that require presence-only data are extremely valuable [17].

Nowadays, the methods that need presence-only data are expanded. One of these methods is Maximum Entropy (MaxEnt) modeling. The MaxEnt model has proven to be highly effective in determining habitat suitability and species distribution because it relies solely on presence data and lacks many of the effects associated with presence-absence analytical methods [22]. The purpose of this study is to model the habitat of Urial (*Ovis orientalis arkal*) in the Samelghan plain (north-eastern Iran) areas with the MaxEnt method. The importance of this region is due to the existence of the Rivi archaeological site and because, based on the samples obtained from zooarcheological studies, the historical distribution of species can be understood. Also, with zooarchaeology and analysis of animal remains, it is possible to reconstruct past habitats [23,24]. Indeed, this study may serve as a pilot project and will give a basic outline for future investigations along with the Tappe Rivi Project that will estimate wildlife versus animal management through ancient times and in comparison to modern fauna.

## Material and methods

### Study area

The Samelghan plain (10 km south of Atrak Valley) is located in Northern Khorasan province, North East of Iran, and exhibits an ancient settlement area of more than 100 ha with four major Tappe ruins. The archaeological remains date back from the Late Bronze and Iron Age throughout the early historic periods of the Achaemenid and Sassanid empires (approx. 1500 BC until 500 AD), plus a succeeding village-like occupation during Early Islamic times [25,26]. The study area covers an area of 1116.6 Km^2^ (Fig. 1), which is located in the geographical position of 37° 21′ to 37° 40′ north latitude and 56° 26′ to 57° 06′ east longitude.

**Fig. 1.**
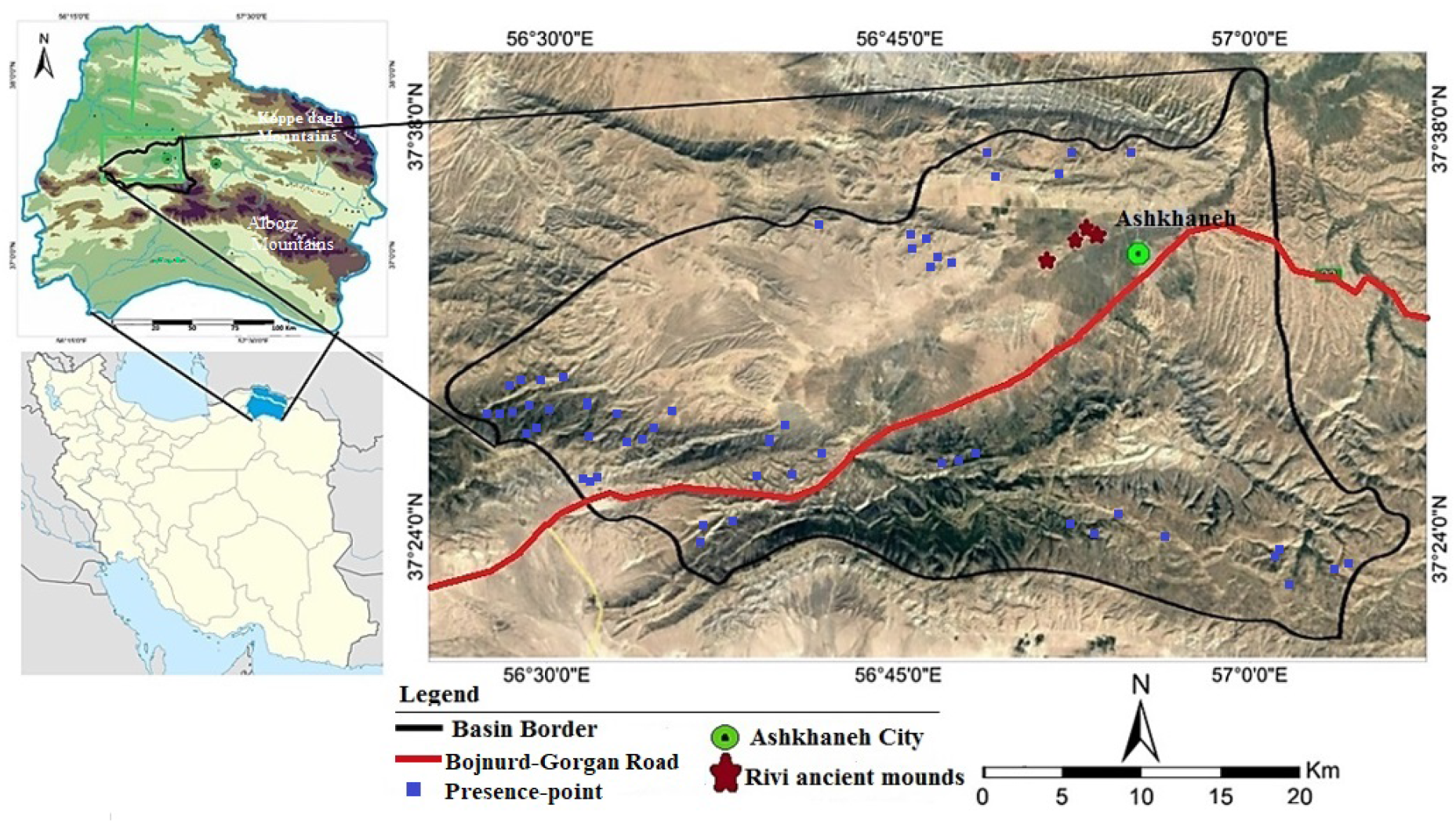
Location of the study area in Iran.

### Species studied

*Ovis orientalis* populations have been decreasing both in size and geographical range in Iran. Poaching, habitat destruction, and competition from livestock have been recognized as the main causes of population decline [27,28]. Urial (*Ovis orientalis arkal*) is a subspecies of *Ovis orientalis* [29] that occurs on rolling hills and gentle mountain slopes in northeast Iran. Urial is currently listed as vulnerable (VU) by IUCN [27].

### Occurrence and Environmental Data

Lack of information about the absence of species complicates the use of conventional environmental modeling tools, because some of these models rely on presence and absence data [8]. For this reason, a modeling technique that does not require absence data was used to identify the environmental factors that explain the distribution of Urial in the Samelghan area. The MaxEnt is a correlative model based on the principle of maximum entropy to predict or infer species occurrence using presence-only data and environmental variables [7,30,31,32,33,34,35]. This algorithm is one of the methods that, despite the small number [30] of presence points have high predictability, and has been widely used by researchers due to time savings and reduced study costs [36,37]. The MaxEnt method uses the presence-only data as a sample and environmental factor classes as environmental variables for modeling in the study area, it also calculates the correlation between the dependent variable and the environment variable [33].

The environmental variables used for modeling include topography and geomorphology, climatology, land use, vegetation, water resources, and human development variables such as villages and roads. Also, class maps of slope percentages and the main aspects were prepared by using DEM. All variables were converted to raster maps after digitization with 30×30 m cell size. After determining the border of the region based on the watershed, all variable were clipped according to the border. In order to collect occurrence records, the distance sampling method [38,39], direct observations, also, information from the province’s environmental experts were obtained. the geographical coordinates of the point were recorded using the Global Positioning System (GPS) as the presence-point, with a total of 53 points (Fig. 1) were obtained for the Urial species in the Samelghan plain.

## Results

The MaxEnt model produces the prediction map, response curves of the variables used,the ROC plot with the AUC (Area Under Curve), and Jackknife’s plot of analysis help researchers to interpret and understand the outcomes of the MaxEnt model. In the MaxEnt method, the map of the potential suitability of the habitat provides a range of suitability for the habitat. The continuous map is shifting between 0 to 1 (Fig. 2) So that if the suitability goes to 1, the habitat for species has higher suitability, and in the same way, it moves to 0, the suitability of the habitat decreases [40].

**Fig. 2.**
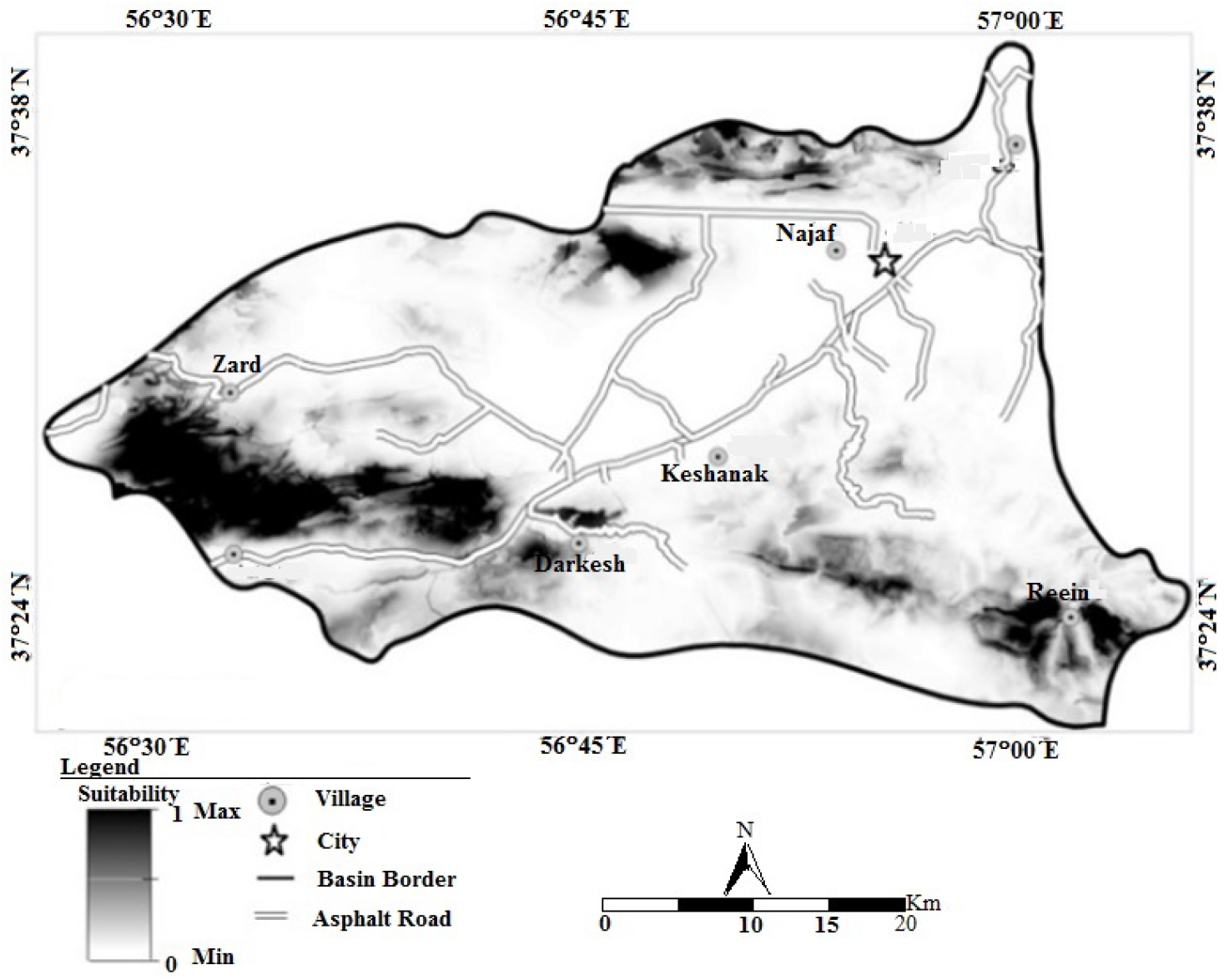
The Habitat continuous map.

In order to use the model in predicting the presence of the species, it is necessary, evaluate and validate the model to determine its accuracy. The area under curve (AUC) Receiver Operating Characteristic (ROC), which equates to the probability of correct detection between presence and absence points by a model, has been widely used as the best evaluation standard in species distribution [41].

The ROC curves evaluate each value of a prediction result as a possible judging threshold [42]. The ROC curve is one of the most common statistical methods widely used in species distribution modeling to evaluate predictive models [7]. The level below the curve (ROC), is the probability of discernment power between the presence and absence data of a model [4343]. The range of AUC values of 0.5 indicates the lowest predictive ability or not different from a randomly selected predictive distribution, and a value of 1.0 indicates perfect model performance [17,44,45, 42]. In this study, the ROC curve for Urial was 0.981, with a standard deviation of 0.001, which indicates the ability to detect very good performance (Fig. 3).

**Fig. 3.**
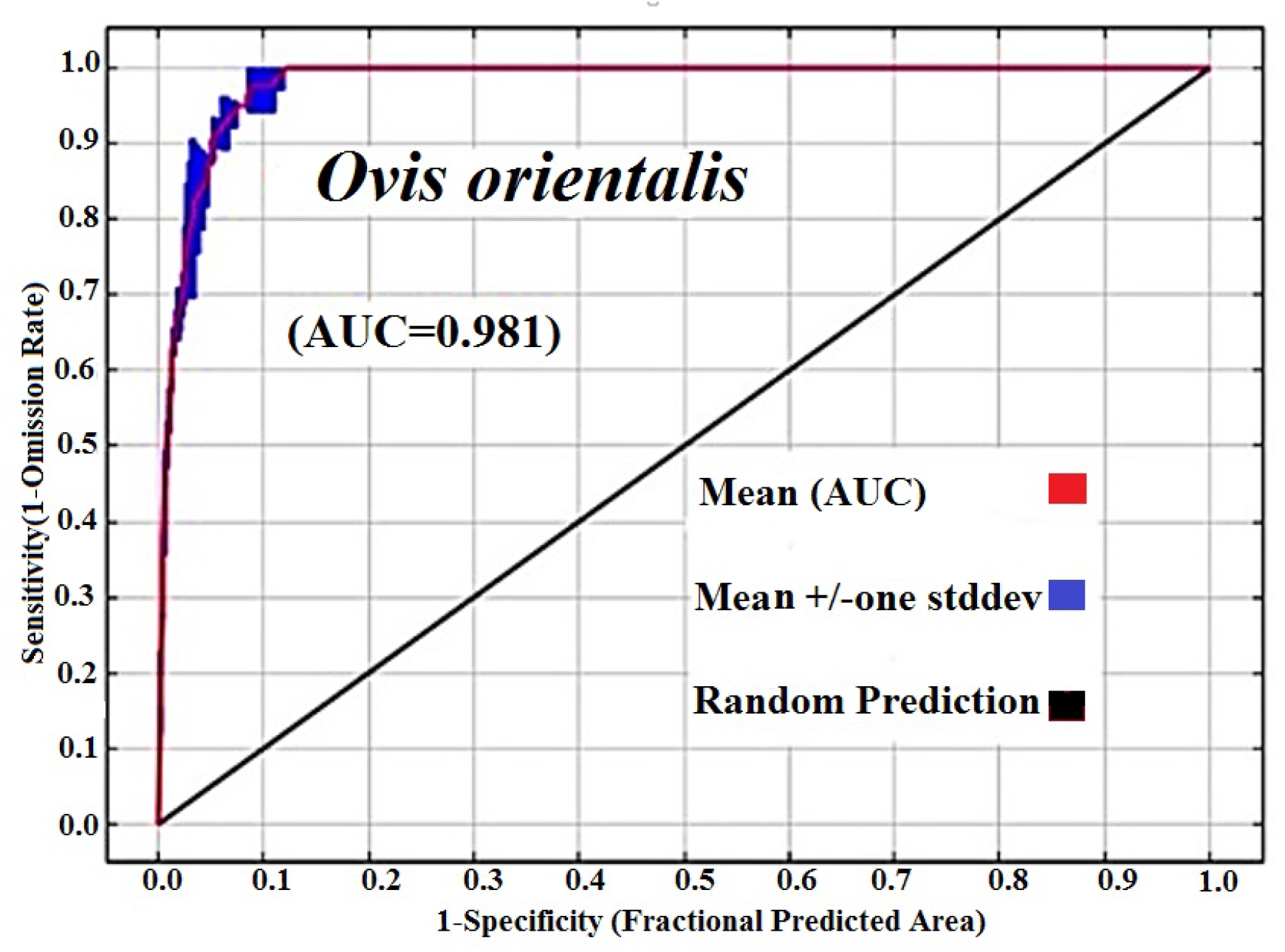
The ROC curve and AUC value.

In the process of modeling species distribution, it is also important to know which variables and to what extent they are involved in predicting species presence. The Jackknife results are displayed in a bar plot to illustrate how the model is run using all input variables, as well as without one variable and using only one variable [46,33]. The Jackknife test provides statistical and accurate estimates of the importance of variables in model prediction. This method removes a variable at runtime and executes the model based on the remaining variables. It then creates a model with each of the dropped variables and finally creates the final model with all the variables participating in the model [47]. The plot shows the importance of variables in three different colors; The blue color indicates how much of the species information is justified when running the model with only one variable, the light green indicates the implementation of the model without the desired variable, and the red color indicates the implementation of the model with all variables [43]. The JackKnife test provides the importance of variables by implementing an effective bootstrap algorithm in the distribution of species [48]. Based on the plot, it can determine which variables alone are most influential in modeling (Fig. 4).

**Fig. 4.**
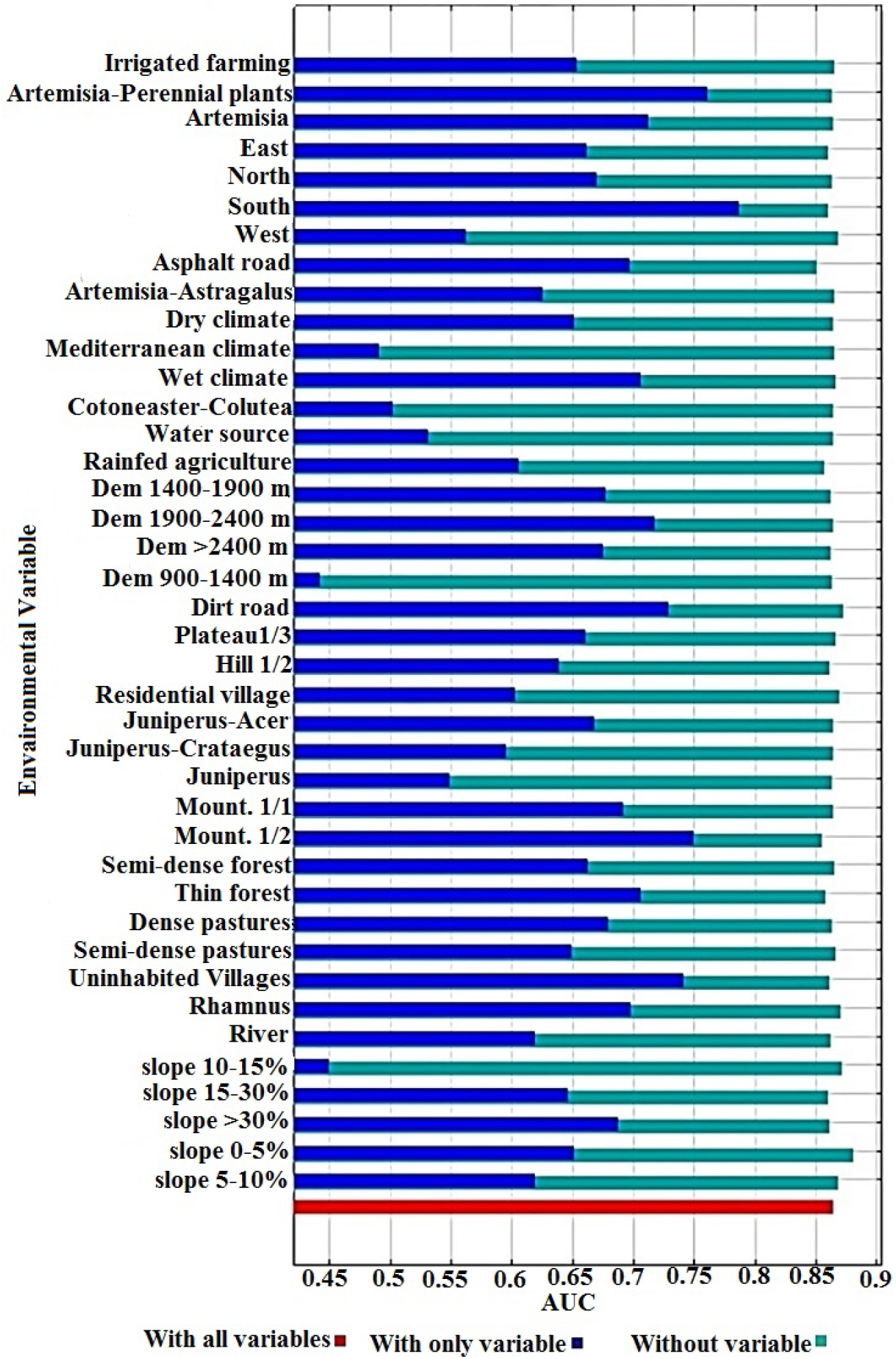
The jackknife plot in determining the importance of variables.

According to our data, the southern aspects are the most effective variable for predicting the distribution of the occurrence data that with remove it, the greatest reduction in the amount of the AUC occurs. These results are used to determine which variables have the greatest influence on probability-presence on the MaxEnt model [33]. Table 1 showed the relative percent values of each of the environmental variables in the distribution of Urial at the surface of the Samelghan plain.

**Table 1.** Percent contribution values of variables used in modeling.

| Variable | Percent contribution | Variable | Percent contribution |
| --- | --- | --- | --- |
| Juniperus-Acer | 14.7 | Eastern Aspects | 1.7 |
| The Slope 0-5% | 12.8 | The slope 5-10% | 1.4 |
| Artemisia | 11.2 | Hill 2/1 | 1.2 |
| Asphalt road | 8.9 | Western Aspects | 1.1 |
| Cotoneaster-Colutea | 6.1 | Plateau 3/1 | 1.1 |
| Artemisia-Perennial plants | 5.5 | Elevation 900-1500 m | 1 |
| Thin forest | 3.1 | River | 1 |
| The slope >30% | 2.4 | Artemisia-Astragalus | 0.9 |
| The slope 10-15 | 2.4 | Rhamnus | 0.7 |
| Mountain 1/2 | 2.2 | Mediterranean climate | 0.6 |
| Elevation >2400 m | 2.1 | Northern Aspects | 0.5 |
| Residential village | 2 | Elevation 1400-1900 m | 0.5 |
| Dense pastures | 2 | Uninhabited Villages | 0.5 |
| Southern Aspects | 2 | Irrigated agriculture | 0.4 |
| Juniperus-Crataegus | 2 | Semi-dense forests | 0.4 |
| Wet climate | 1.9 | Rainfed agriculture | 0.3 |
| Semi-dense pastures | 1.9 | Juniperus | 0.1 |
| Dirt road | 1.7 |  |  |

Created map for species was entered into the ArcGIS10.3 and according to the threshold value obtained from the model (0.0513) for Urial, the habitat was divided into a binary map (suitable/unsitable areas) (Fig. 5). The final habitat suitability map has been prepared from the results and interpretation of the Jackknife plot, the response curve, and the ROC curve based on the presence-only data of species in the region.

**Fig. 5.**
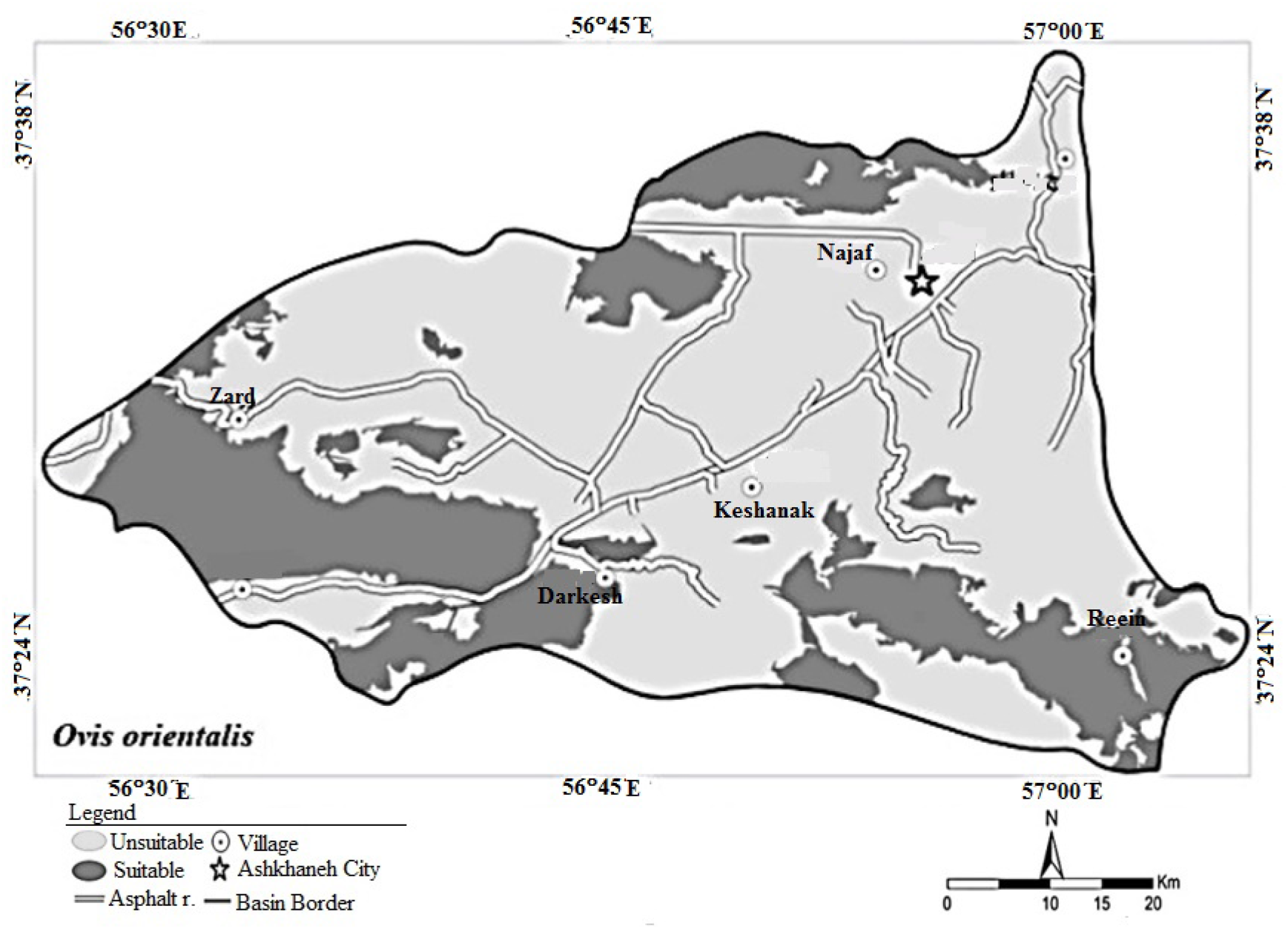
The habitat classification map.

## Discussion

Habitat suitability models are useful tools for a variety of wildlife management objectives. Distributions of wildlife species can be predicted for geographical areas that have not been extensively surveyed [49]. Habitat models are also useful for predicting areas that may not currently be used by wildlife species [50]. The basis of these models’ work is to minimize the relationship between species distribution and biotic and abiotic environments [51].

The MaxEnt identifies important variables in determining suitable/unsuitable habitats and the importance of each variable in increasing or decreasing the likelihood of species presence the space. The AUC value obtained was 0.981, therefore, this model provide new information about the potential distribution of the species with fairly good accuracy.

The most important variables are obtained based on the “percent contribution values” and “Jackknife test” include; Juniperus-Acer, the areas with a slope of 0-5%, Artemisia-Perennial Plants, mountain 1/2 (high mountains with deep valleys consisting of limestone, metamorphic and igneous rocks), southern aspects, and asphalt road. These variables have the most impact on modeling compared to other variables used in the model, that is by eliminating each of these variables, the most change will be in the chart and finally the most change in the final model.

The slope variable of 0-5% is one of the variables that were important in Maxent’s results. The species response curve for this variable showed that this factor was effective in the absence of species and reduced suitability. In other words, areas with higher slopes are more important in the presence of the species and habitat suitability.As shownin figure 6, steep slopesare so important in the presenceof the species that the suitability of the habitat will begreatly reduced by increasing the distance. In studies that have been shown in other areas for Urial species, high slopes classes are more important [48]. The asphalt road variable is important in habitat modeling of the study area because of its high abundance in the basin and species habitat. This variable causes the habitat fragmentation. The probability of presence is highrt in habitats that are far from this variable, so that with increasing distance from this variable, the suitability of habitat increases as well. In the present study, the results showed that the species differently response to each of the environmental variables. Thus, some variables contribute more to modeling, That is, they are very effective in modeling the presence of the species. This function can have a variety of causes, including the presence of a competing species, the hunters, the food sources, and many other factors that affect the presence of the species.

**Fig. 6.**
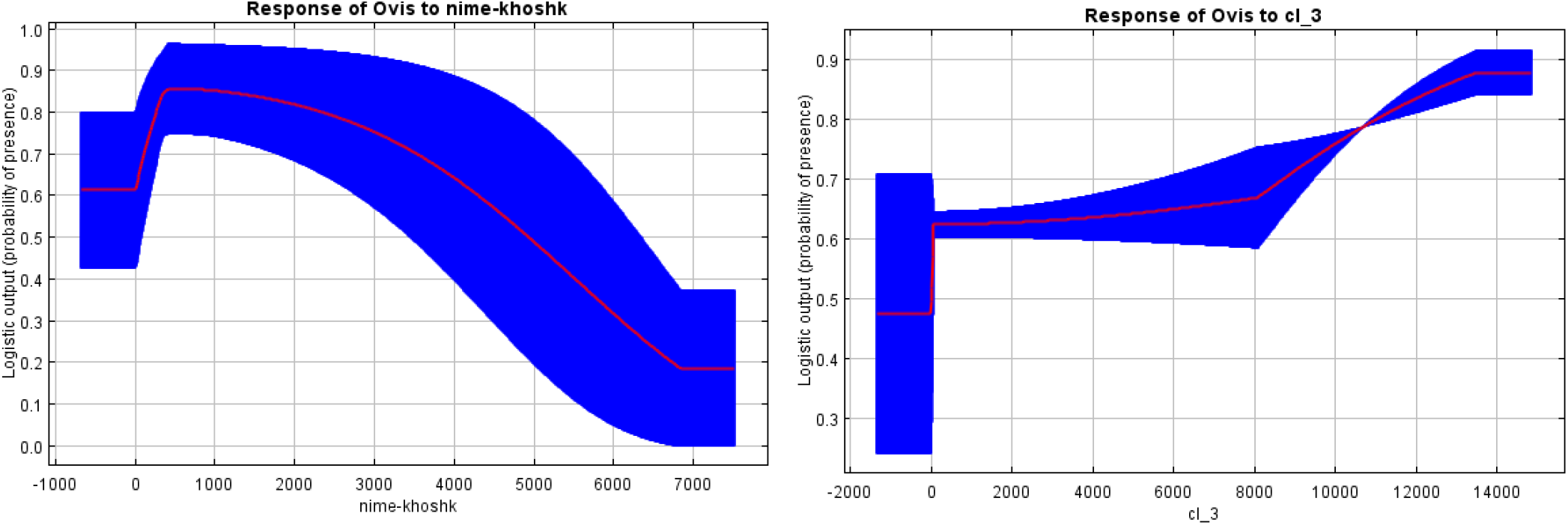
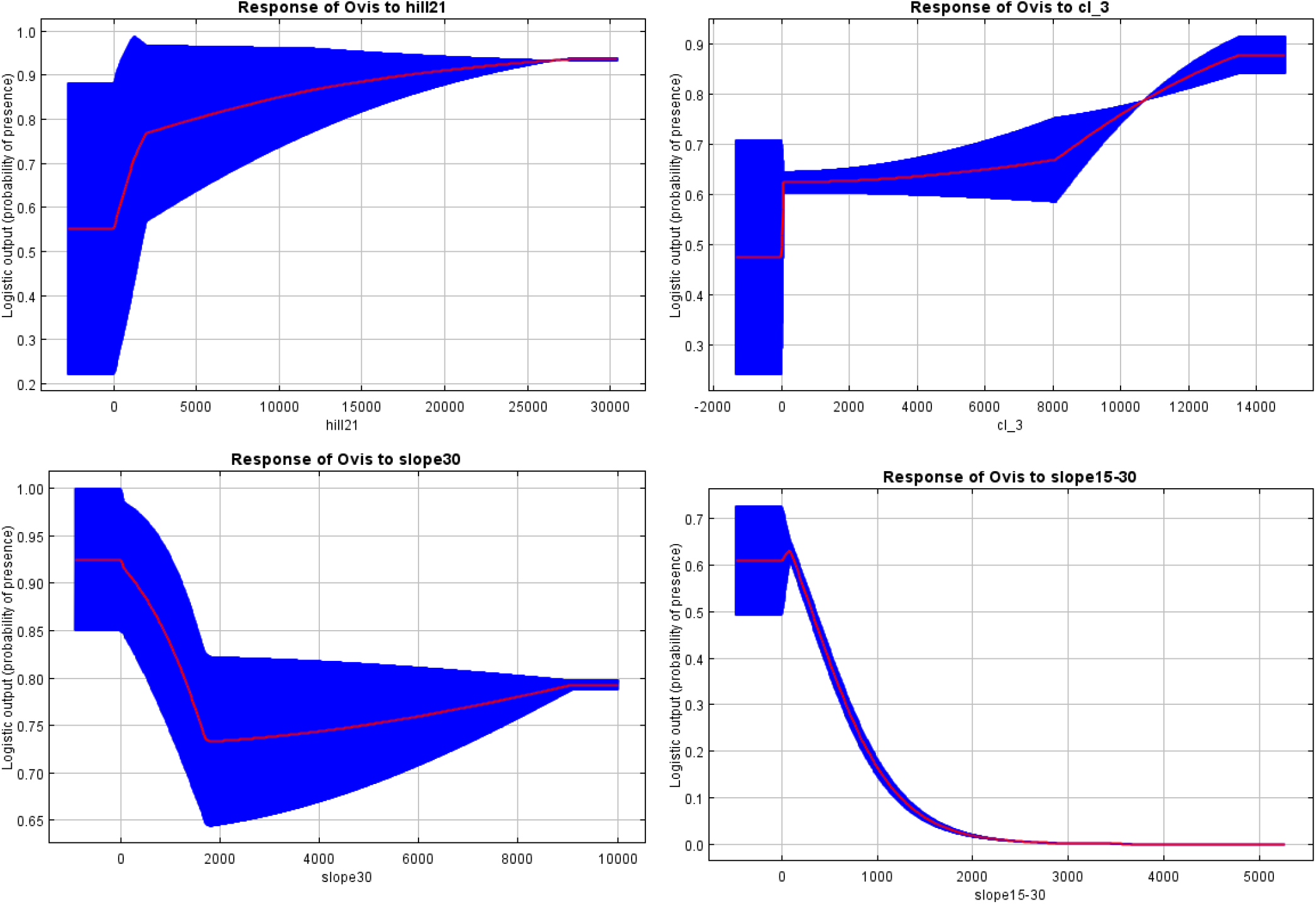
The response curves of Urial to environmental variables.

Our results show that environmental variables cause significant changes in the suitability models for the Urial and based on these changes, management decisions can be made to protect species. Indeed, it is suggested that:

- since the area was not under cultivation, other species should be identified and examined, if possible, to define the area as a “No-hunting area”.
- along with future results of the zoo-archaeology for the ancient periods, one may also ancient wildlife habitats modeling and possible changes as well as the impact of human-animal management.

Habitat suitability map in the Samelghan plain shows that habitats are important for Urial that are outside residential areas and human activities. However, some areas, such as roads and parts of agricultural lands are in the suitable habitat for the species, it is because of the expansion of roads and the development of human activities in the study area.

Finally, Maxent produces the final map based on the effects of all environmental variables that effect increasing and decreasing suitability. Suitable habitat indicates the importance and interaction of all the environmental classes used in modeling. This study showed that approximately 28% (313 Km^2^) of the Samelghan plain is a suitable habitat for the Urial. The habitat’s predicted in the areas of the region, which has at least overlap with human activity, also, least conflict than other parts of the region. These areas include a habitat that has better quality in terms of nutrition and shelter. According to the obtained map, it can be understood to what extent human activities have been effective in fragmenting the habitat of Urial . Areas suitable for Urial are mostly extended in the highlands of the study area, including hills and mountains.

